# Electroencephalographic markers of brain development during sevoflurane anesthesia in children aged 0 to 3 years old

**DOI:** 10.1101/165902

**Authors:** Laura Cornelissen, Seong-Eun Kim, Johanna M. Lee, Emery N. Brown, Patrick L. Purdon, Charles B. Berde

## Abstract

The general anesthetic sevoflurane acts on GABAergic-interneurons to generate stereotyped oscillations that relate fundamentally to neural circuit architecture. Each year, millions of children require general anesthesia, providing an experiment of nature that allows characterization of the developmental trajectory of GABAergic-inhibitory circuits in the human brain. We used multichannel electroencephalograph recordings in 91 children 0-40 months old. We mapped spatial power and coherence across the cortex. During sevoflurane exposure: (1) slow-delta oscillations are present in all ages; (2) theta and alpha oscillations emerge around 4 months; (3) alpha oscillations increase in power from 4 to 10 months; (4) frontal alpha oscillation predominance emerge at ~6 months; (5) frontal slow oscillations are coherent in the first 6 months of age only; and (6) frontal alpha oscillations become coherent around 10 months and persist in older ages. Our results suggest key developmental milestones are visible in the functional activity of sevoflurane-stimulated GABAergic circuits.

## Introduction

Brain development spans more than two decades in humans from embryo to young adulthood. Gamma-aminobutyric acid (GABA) is the major inhibitory neurotransmitter in the brain and plays a critical role in driving the assembly and fine-tuning of neural circuits throughout this protracted period; disruptions of these processes are thought to underlie many developmental disorders including autism (Hensch, 2005; Marín, 2016). Despite the importance of inhibition in the brain, trajectories for development of these GABAergic inhibitory brain circuits have been characterized in animal models, but less so in humans because of the difficulty in measuring circuit activity.

Each year millions of children are administered general anesthesia for surgery (DeFrances & Hall, 2007; Rabbitts et al., 2010). Volatile anesthetic drugs bind to multiple targets at the brain and spinal cord to exert their physiological and functional effects (Brown et al, 2011; Purdon et al., 2015). Sevoflurane is one of the most commonly-used vapor anesthetics in children and is used for its rapid induction, emergence and recovery profile. Sevoflurane acts primarily to potentiate GABA_A_ inhibition (Hemmings et al., 2005), providing an experiment of nature that makes it possible to characterize how the brain responds to a strong GABA stimulus as a function of age.

Clinical studies using non-invasive brain monitoring show general anesthetics and hypnotic agents such as sevoflurane, propofol, ketamine and dexmedetomidine produce stereotyped electroencephalograph (EEG) oscillations that relate fundamentally to neural circuit architecture and function (Brown et al., 2010; Purdon et al., 2015). In adults under sevoflurane or propofol anesthesia, structured EEG oscillations with specific spatial organization are present (Akeju, et al., 2014; Cimenser et al., 2011; Pavone et al., 2017; Purdon et al., 2015; Purdon et al., 2013). For example, under propofol anesthesia, specific EEG patterns consisting of incoherent slow oscillations distributed across the entire head and strongly coherent alpha oscillations located across only the front of the head are observed (Purdon et al., 2013). Thalamic, cortical and thalamocortical relay cells are thought to play an important role in the generation of the oscillation brain rhythms. Theoretical modelling approaches based on propofol (which has comparable EEG properties to sevoflurane) suggest that propofol increases GABA_A_ conductance, which in turn facilitates activity of a highly coherent alpha oscillation loop between the thalamus and frontal-cortex (Ching et al., 2010; Vijayan et al., 2013).

Few studies detailing the neural circuit activity of sevoflurane in children have been described. Preliminary studies suggest that the neurophysiologic response to sevoflurane are different from that of adults (Akeju et al., 2015), and change as a function of age (Akeju et al., 2015; Cornelissen et al., 2015; Davidson et al., 2008; Hayashi et al., 2012; Sury et al., 2014). However, these studies lack detailed characterization of anesthesia-associated neural circuit activity in relation to spectral and coherence properties, and spatial distribution of neural oscillations at specific ages where critical periods of development take place.

Given that the brains of children in different age groups are at different developmental stages due to changes in GABAergic circuitry, we investigated whether there are systematic changes in sevoflurane-induced EEG dynamics that mirror these different developmental states. We therefore analyzed in detail how sevoflurane-induced brain oscillations change across the first three years of life. We used multichannel scalp electroencephalograph (EEG) recordings in children undergoing sevoflurane general anesthesia for elective surgery, to determine transitions in EEG power spectra and coherence across early infancy (birth to 3 years old). We mapped spatial power and coherence over frontal, central, parietal, temporal and occipital cortices. Our results suggest key developmental milestones in the assembly and maintenance of GABAergic circuits in the human brain.

## Results

Continuous multichannel EEG recordings collected during maintenance of a surgical state of anesthesia (MOSSA) in 91 subjects 0 to 3 years of age (0 to 40 months old) are reported. Subject demographics and clinical characteristics are provided in ***Table 1***. Details concerning study design are given in the ‘*Materials and Methods*’ and ***Figure 1***.

**Table 1.**
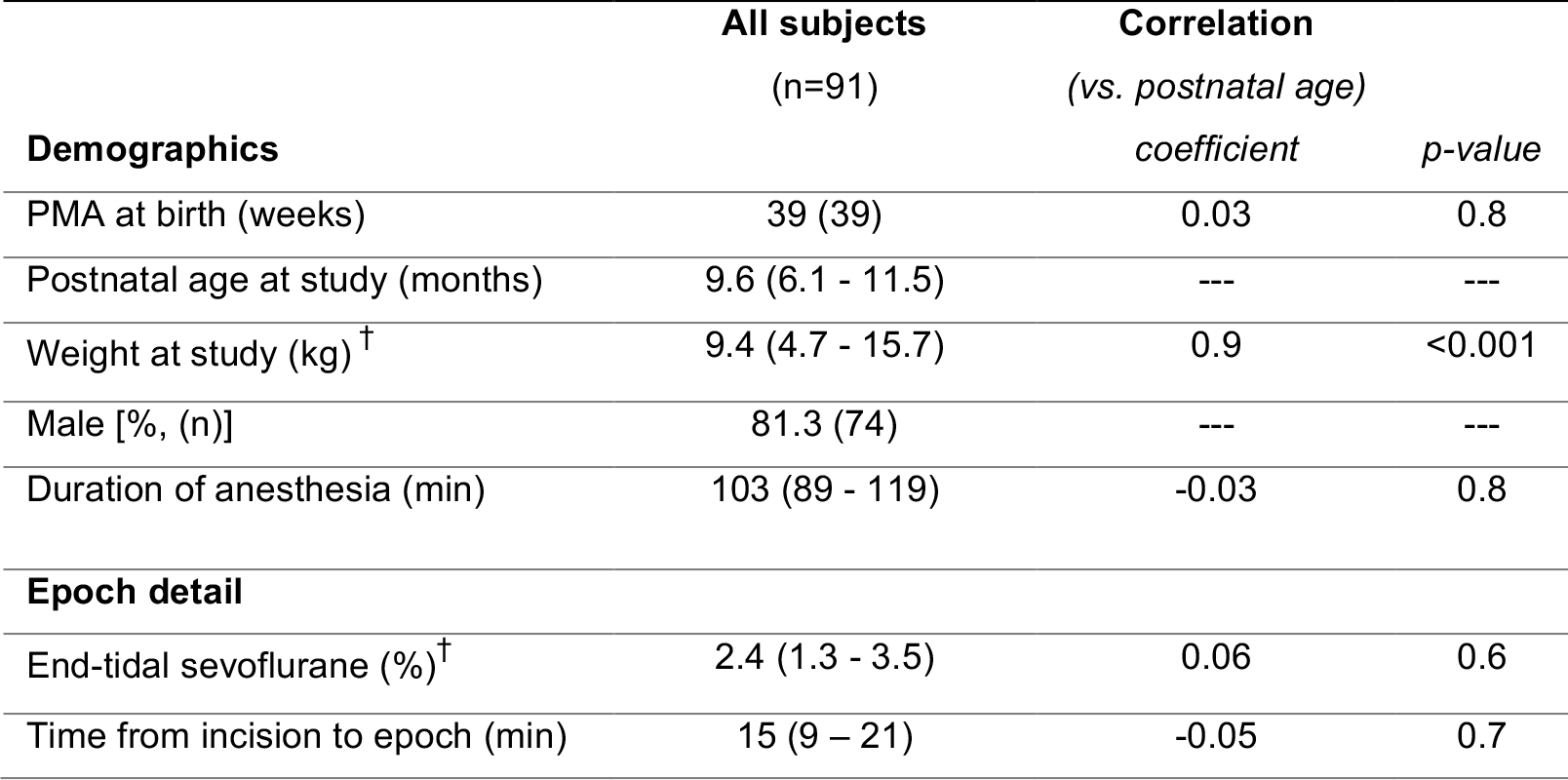
Subject demographics and clinical characteristics Data given as median with 95% Confidence Interval (CI) limit in parentheses unless otherwise stated. Pearson’s product moment or Spearman’s rank order coefficient were computed to assess the relationship between postnatal age at the study and each continuous column variable; *r* (with 89 degrees of freedom) and *p* are reported, p<0.05 was considered statistically significant. ^**†**^Data given as mean with 95% CI limit in parentheses, with corresponding Spearman’s *r* and p-value.

|  | **All subjects**<br>(n=91) | **Correlation**<br>(vs. *postnatal age*) |  |
| --- | --- | --- | --- |
| **Demographics** |  | *coefficient* | *p-value* |
| PMA at birth (weeks) | 39 (39) | 0.03 | 0.8 |
| Postnatal age at study (months) | 9.6 (6.1 - 11.5) | --- | --- |
| Weight at study (kg)<sup>†</sup> | 9.4 (4.7 - 15.7) | 0.9 | <0.001 |
| Male [%, (n)] | 81.3 (74) | --- | --- |
| Duration of anesthesia (min) | 103 (89 - 119) | -0.03 | 0.8 |
| **Epoch detail** |  |  |  |
| End-tidal sevoflurane (%)<sup>†</sup> | 2.4 (1.3 - 3.5) | 0.06 | 0.6 |
| Time from incision to epoch (min) | 15 (9 - 21) | -0.05 | 0.7 |

**Figure 1.**
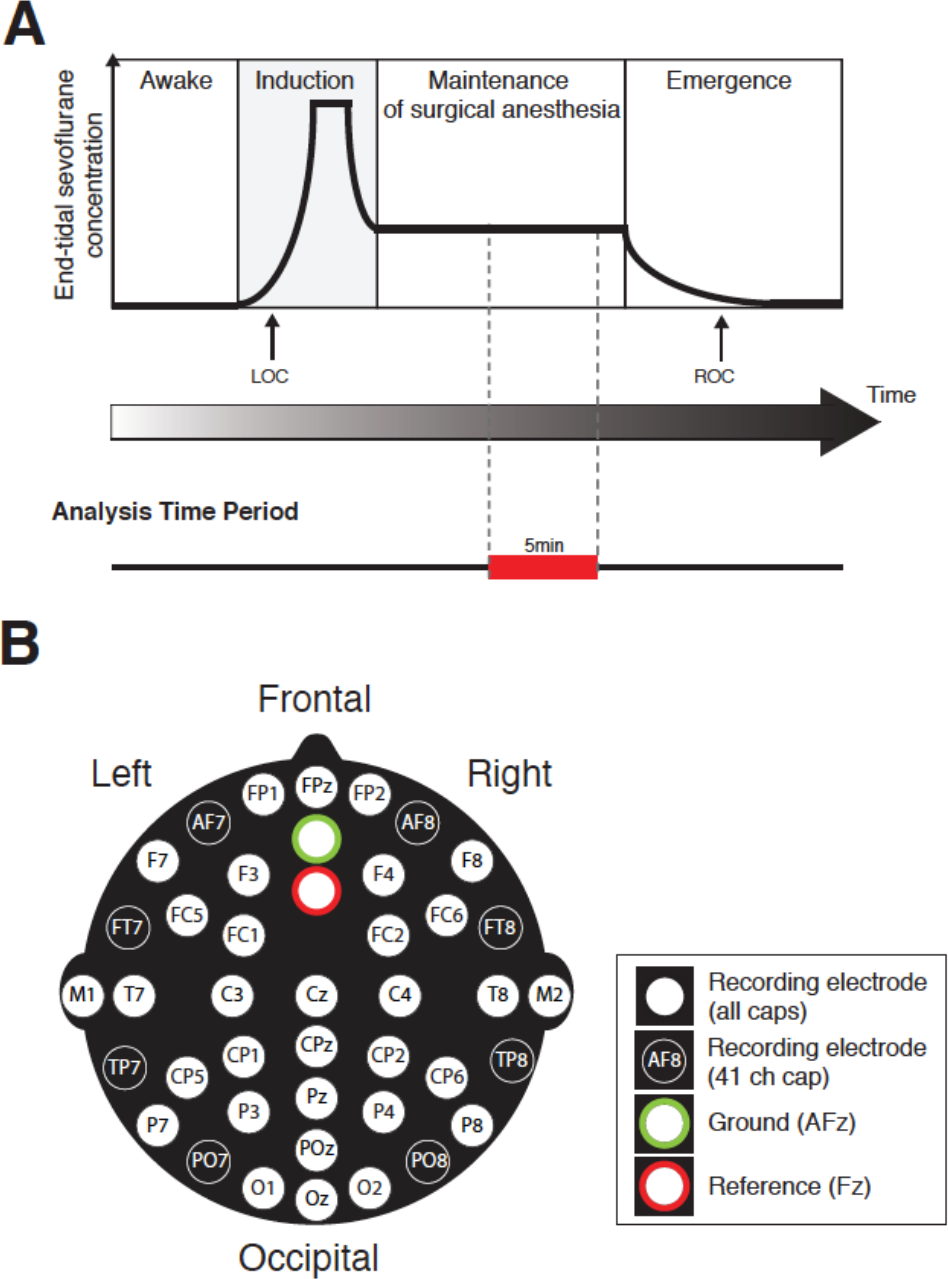
Study design. Study timeline (A): schematic time-course of end-tidal sevoflurane concentration during the awake phase, induction, maintenance of surgical anesthesia, and emergence phases of general anesthesia. Electroencephalogram (EEG) data from individual recording electrodes were analyzed post-hoc during maintenance of surgical anesthesia using an epoch size of 5-minutes (shown in red). (B) EEG montage used, 33-channel EEG caps were used in patients weighing <10kg, and 41 channel (ch) EEG caps were used for patients weighing ≥ 10kg (modified international 10-20 electrode placement system).

### Age-varying changes in power spectral properties

#### Slow and delta oscillations are present across all ages; theta and alpha emerge around 4 months of age

Frontal power spectra analysis in individual infants at electrode F7 show that slow (0.1-1 Hz) and delta power (1-4 Hz) were present at all ages during maintenance of sevoflurane anesthesia (***Figure 2A***). At the youngest ages –1 and 3 months old, power in the 4-40 Hz frequency range was minimal. At 5 months old, power in the theta (4-8 Hz) and alpha (8-12 Hz) frequency ranges began to emerge. At 9, 11 and 16 months, higher frequencies in the low beta (13-17 Hz) ranges appear. At 19 and 37 months, the alpha frequency range began to increase in power compared to other frequencies above 4 Hz. Based on these observations, subjects were subsequently separated into groups according to age: 0-3 months (n=17), 4-6 months (n=23), 7-9 months (n=7), 10-14 months (n=18), 15-17 months (n=7), and 18-40 months (n=19) for subsequent grouped-analysis (***Figure 2 – figure supplement 1***).

**Figure 2.**
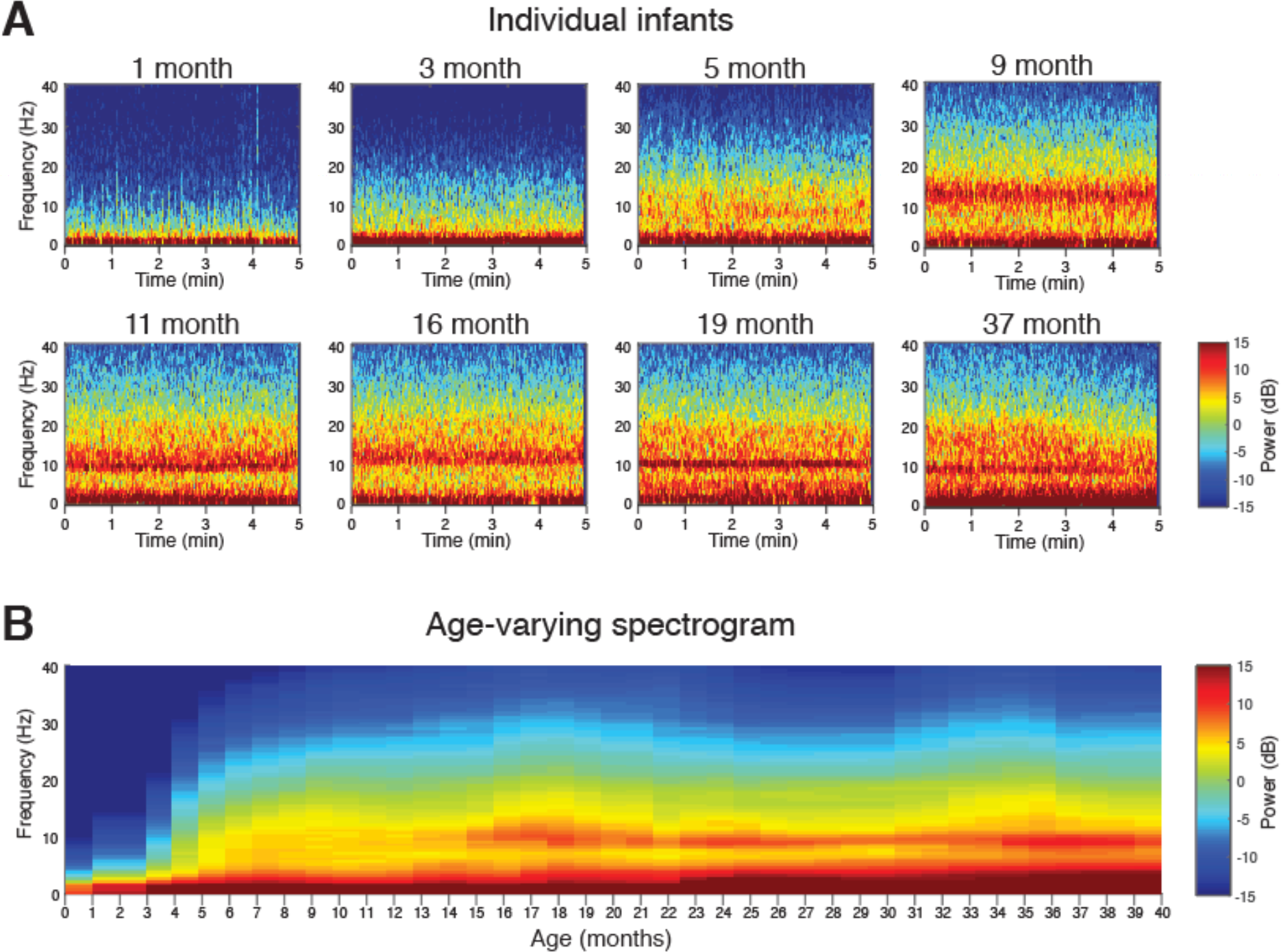
Frontal spectrograms during sevoflurane-maintained surgical anesthesia in children from 0 to 3 yrs. Frontal EEG spectral power for frequencies from 0.1 to 40Hz during a 5-min period of surgical anesthesia. Frontal spectrogram for (A) individual subjects at 1, 3, 5, 9, 11, 16, 19 and 37 months of age. Age-varying spectral changes in (B) frontal spectrogram. F7 used with nearest neighbor Laplacian referencing.

The age-varying frontal spectrogram (time-varying spectra) computed at F7 shows the distribution of slow-delta, theta, alpha, beta and gamma (25-40 Hz) oscillations by age (***Figure 2B***). Slow and delta were dominant frequencies across all ages. Theta and alpha oscillations were more prominent in subjects aged 4 months and older. Beta and gamma oscillations remained less prominent of all frequencies. Age-grouped analysis of frontal group-median power spectra computed at F7 show the evolution of alpha oscillations with age (***Figure 2 – figure supplement 1***).

Slow, delta and alpha oscillations were the dominant frequency components observed across age. Peak frontal power was evaluated for slow, delta and alpha frequencies across age. Frontal slow and frontal delta oscillations were present from birth and steadily increased with age (***Figure 3A & B***). Frontal alpha oscillations emerged at approximately 3 to 4 months and steadily increased until peaking at 10 months of age (***Figure 3C***). Alpha oscillations remained at a consistent power level from 10 months to 3 yrs (~6dB).

**Figure 3.**
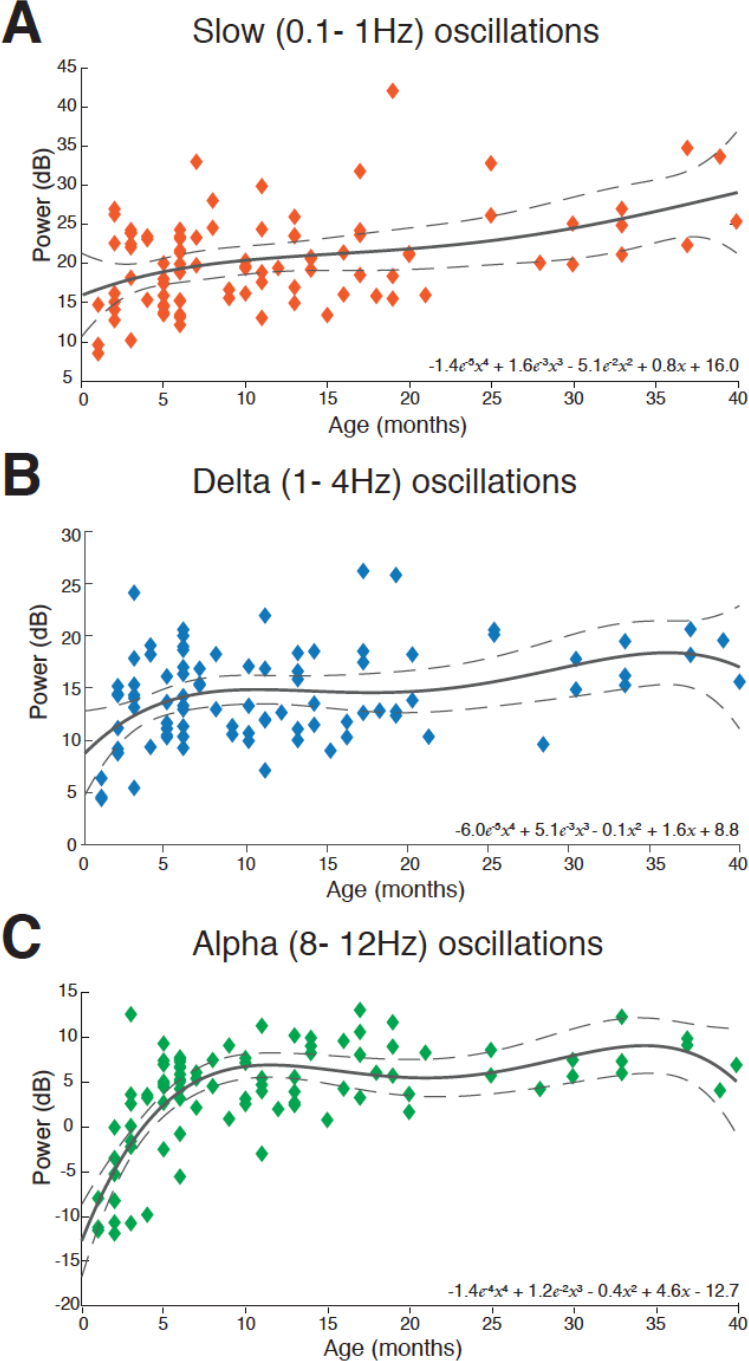
Trends in frontal slow, delta and alpha power with age from 0 to 3 yrs. Peak EEG power in the (A) slow (0.1 to 1Hz), (B) delta (0.1 to 4Hz), and (C) alpha (8-12Hz) frequency bands for each subject during sevoflurane-maintained surgical anesthesia, plotted as a function of age. Delta oscillations exhibited a small, steady increase in power with age. Alpha oscillations increased in power in the early postnatal age, peaking at approximately 10 months and remaining at a sustained level with age. Solid line represents a fourth-degree polynomial regression model describing the relationship between age and EEG power, with equation written in inset; dashed lines represent the 95^th^ confidence boundaries of this regression model. F7 electrode presented using nearest neighbor Laplacian referencing. Epoch size of 5 minutes is used.

#### Frontal alpha predominance emerges around 7 to 9 months of age

Topographic plots were computed for slow, delta, theta, alpha, beta and gamma frequency bands in each age-group (***Figure 4***). Spatial distribution of EEG spectral power for each electrode location in individual infants are shown in ***Figure 4 – figure supplement 1***. For all age-groups, slow, delta and theta activity were generally widely distributed across the scalp. Alpha activity was present to a greater degree from around 4 months, and was largely regionalized to the frontal cortex from around 7 to 9 months of age. Beta and gamma activity remained negligible across all ages.

**Figure 4.**
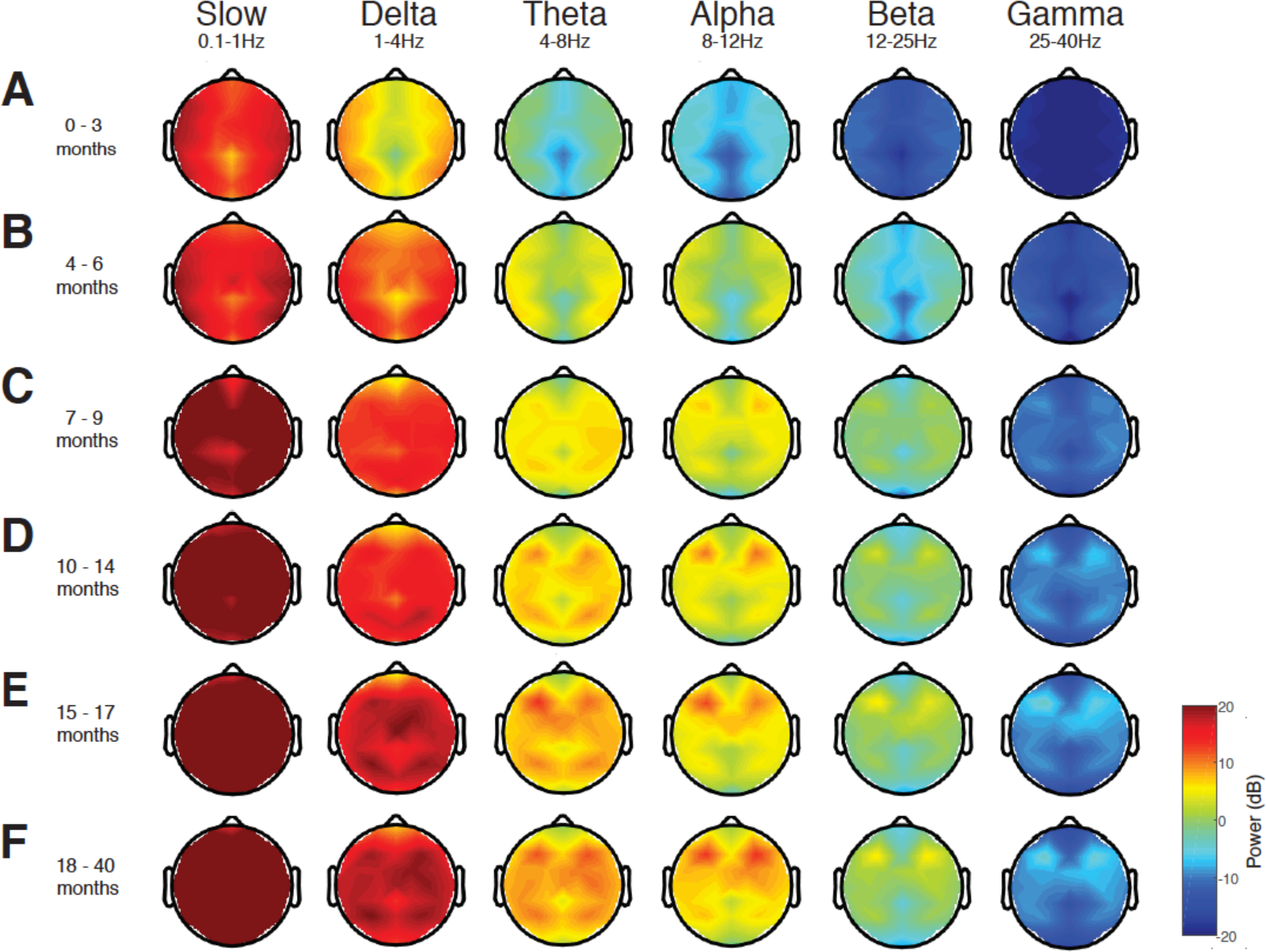
Topographic EEG maps of spectral power for distinct frequency bands during sevoflurane-maintained surgical anesthesia in children 0 to 3 yrs. Topographic EEG maps detailing group-averaged power for each EEG frequency band in infants aged (A) 0-3 months (n=17), (B) 4-6 months (n=23), (C) 7-9 months (n=7), (D) 10-14 months (n=18), and toddlers aged (E) 15-17 months (n=7), and (F) 18-40 months (n=19). Slow wave, delta and theta activity is distributed across the scalp in all age groups. Alpha activity is present to a greater degree from around 4 months, and is largely regionalized to the frontal cortex from around 7 months of age. Beta and gamma oscillations remain negligible across all ages. Nearest neighbor Laplacian referencing scheme with an epoch size of 5 minutes is used.

For infants 0 to 3 months, spectral power along the midline (specifically central-parietal locations) was lower across all frequencies. For subjects older than 3 months, spectral power at the central-parietal, and occipital midline was lower in the theta and alpha ranges. In all ages, slow and delta oscillations were broadly distributed across the scalp. Theta oscillations were also broadly distributed across the scalp, and became more pronounced at around 10 to 14 months of age. For subjects 4 to 6 months, alpha oscillations were broadly distributed across the scalp with lower power in the central-parietal and occipital midline. For subjects older than 6 months, peak alpha power was located over frontal regions, and expanded to include the central area with increasing age.

Frontal predominance of alpha power was evaluated by assessing power spectra differences between frontal (F3) and occipital (O1) electrodes across ages (***Figure 5***). Frontal electrodes (F3, F4) were identified as the locations with the greatest total power in the alpha frequency band from the topographic mapping, and were therefore used in this analysis. Frontal alpha oscillations were consistently greater in power compared to occipital oscillations from around 8 months of age and older. Slow and delta oscillations remained uniformly distributed over the frontal and occipital cortices. These data indicate that adult-like patterns of activity, anteriorizaton of alpha oscillations, are detectable from late infancy onwards.

**Figure 5.**
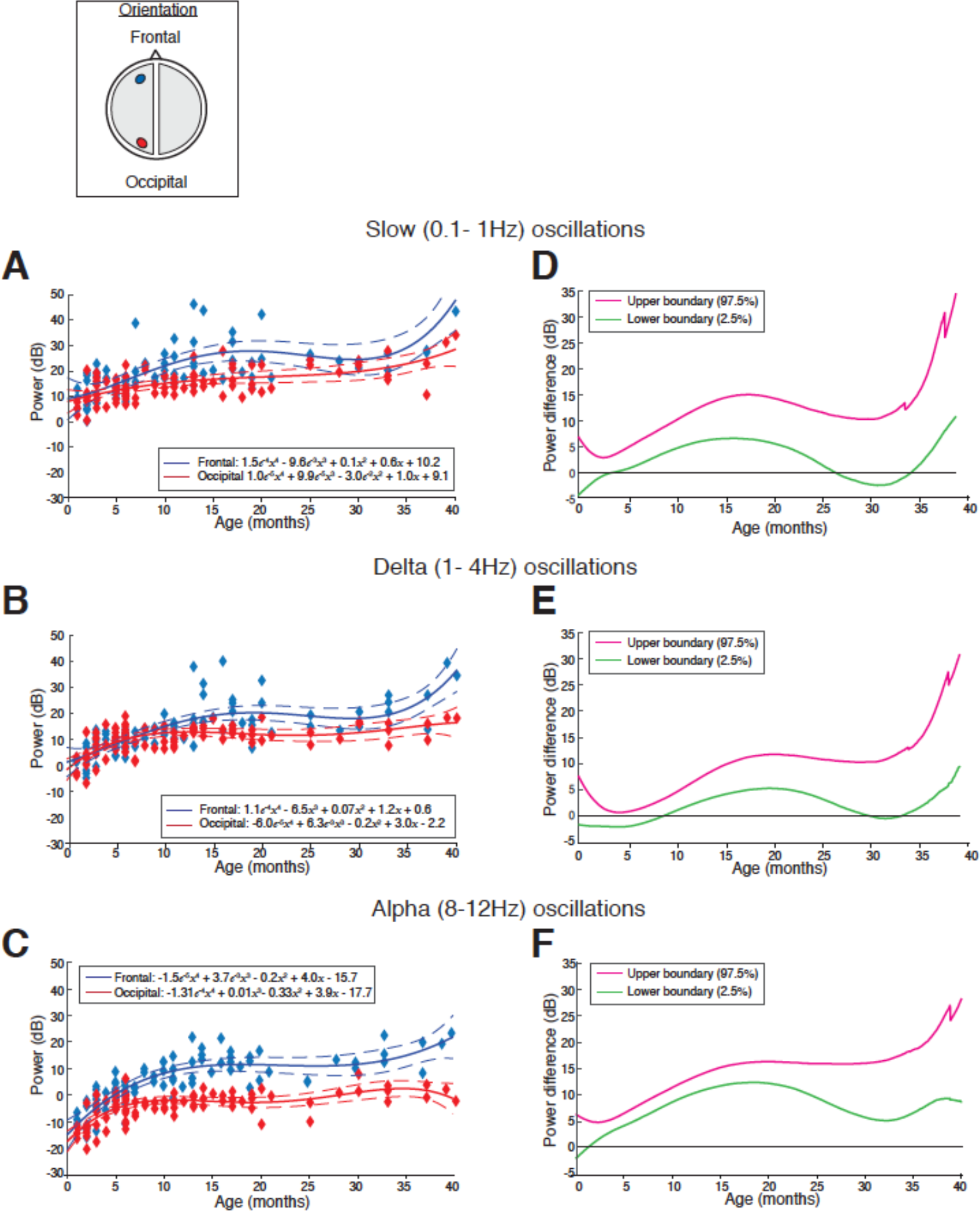
Frontal and occipital power distribution at slow, delta and alpha frequencies during sevoflurane-maintained surgical anesthesia in children from 0 to 3 yrs. Peak frontal and occipital EEG power in the (A) slow (0.1 -1Hz), (B) delta (1-4Hz), and (C) alpha (8-12Hz) frequency bands for each subject are plotted as a function of age. Solid line represents fourth degree polynomial regression model describing the relationship between age and EEG power (blue line, frontal power; red line, occipital power; dashed lines, 95^th^ percentile confidence boundaries); inset schematic indicates the location of the frontal and occipital channels. Differences in power between frontal and occipital channels in the (D) slow (0.1 -1Hz), (E) delta (1-4Hz), and (F) alpha (8-12Hz) frequency bands, are presented with 95% CI from bootstrap analysis (pink line, 97.5^th^ percentile; green line, 2.5^th^ percentile). Slow and delta oscillations exhibit a small and significant increase in frontal power compared to occipital power from mid-late infancy until ~26 to 30 months of age. Alpha oscillations emerge around 3-4 months of age, and exhibit a significant and sustained frontal predominance of power that begins to emerge at 8 months of age, and peaks at ~15 to 20 months of age. These data indicate that adult-like patterns of activity, are detectable from infancy. F3 and O1 electrodes presented using nearest neighbor Laplacian referencing. Epoch size of 5 minutes is used.

### Age-varying changes in coherence properties

#### Frontal slow-delta coherence is present from birth to 3 months of age; frontal alpha coherence emerges at 10 months

Frontal coherence - the level of local coordinated activity in the frontal regions of the scalp - during sevoflurane-maintained surgical anesthesia was analyzed by computing the coherogram (time-varying coherence) and coherence between the left (F7-FP1) and right (F8-Fp2) frontal electrodes. Frontal coherence analysis in individual infants showed trends across age. At the youngest ages –2 and 3 months old, slow and delta oscillations were highly coherent (>0.6). At 5 months were slow and delta oscillations were less coherent, and at 9 months were incoherent (uncoordinated). At 16 months old, alpha oscillations began to exhibit coherence. At 19 and 37 months old, slow and delta oscillations remained incoherent, and in contrast, alpha oscillations were highly coherent (***Figure 6A***).

**Figure 6.**
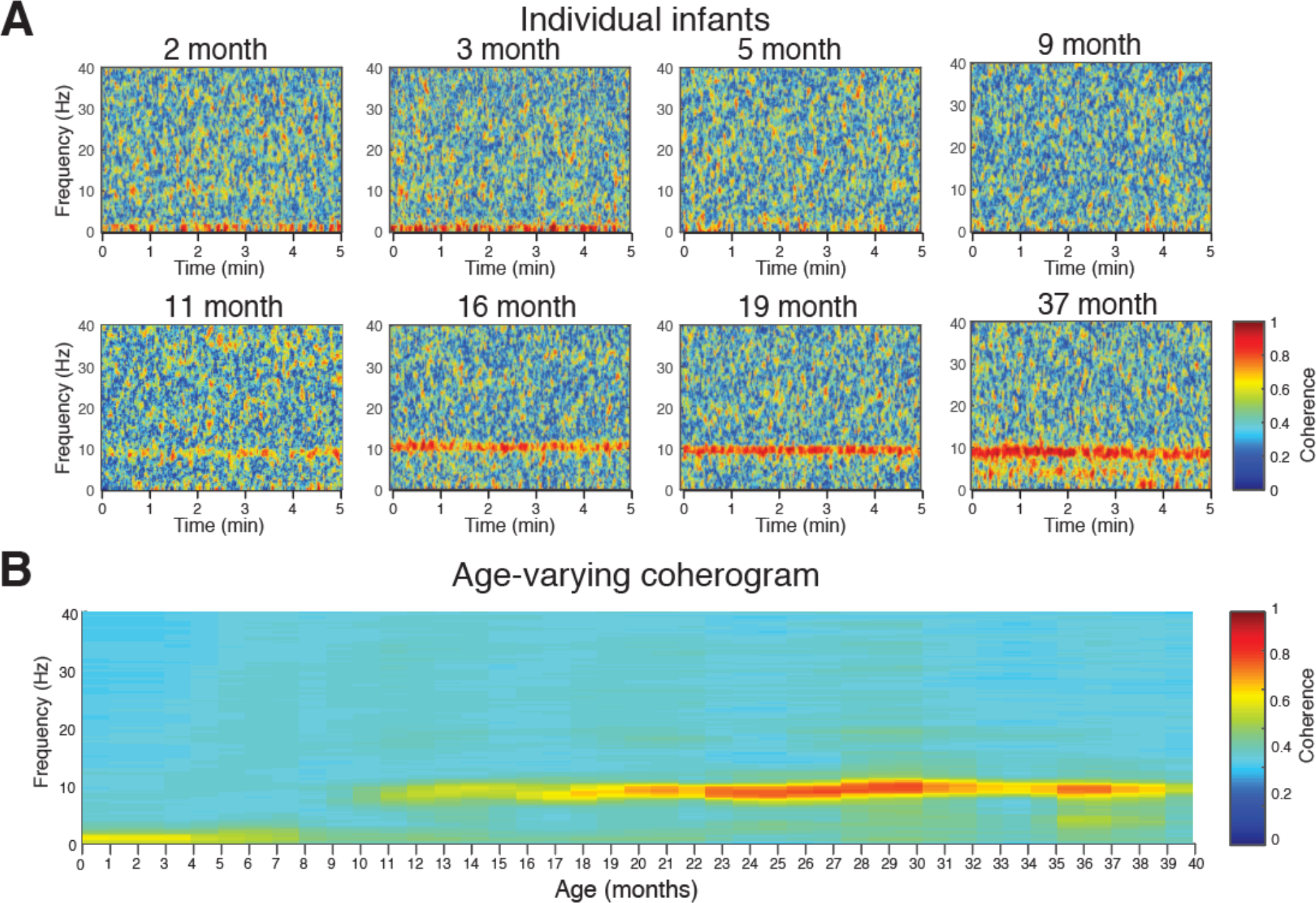
Frontal coherograms during sevoflurane-maintained surgical anesthesia in children from 0 to 3 yrs. Frontal EEG coherence power for frequencies from 0.1 to 40Hz during a 5-min period of surgical anesthesia. Frontal coherogram for (A) individual subjects at 2, 3, 5, 9, 11, 16, 19 and 37 months of age. Age-varying coherence changes in (B) frontal coherogram. Slow (0.1-1Hz), and delta (1-4Hz) oscillations are coherent in subjects from birth until around 8 months old. Alpha coherence appears to emerge around 10 months of age and gradually increases in 20 months where it persists to 3 years of age. Left frontal (F7-Fp1) to right frontal (F8-Fp2) coherence is shown. Nearest neighbor Laplacian referencing and an epoch size of 5 minutes is used.

The age-varying frontal coherogram shows the distribution of slow-delta, theta, alpha, beta and gamma oscillations (***Figure 6B***). Slow and delta were dominant coherent frequencies from birth until around 8 months of age, with the peak coherence in the first three months (~0.6 – 0.7). Slow and delta oscillations were incoherent in children from 9 months of age and older. Despite alpha oscillations being present in the spectrogram from 3 to 4 months of age, alpha oscillations were incoherent at this age and only became coherent at around 10 months of age (at very low levels and with variability across subjects). Gradually alpha oscillation coherence increased until 20 months where it persisted through to 3 years of age. These results suggest that infants in the first 8 months of life have coordinated local frontal slow and delta oscillations, while from 10 months of age the fine-tuning of coordinated alpha oscillations begins to emerge. This is similar to adults who show a strong frontal coherence in the alpha band during sevoflurane general anesthesia (Akeju et al., 2014). Age-grouped analysis of frontal group-median coherence computed show the maturation of alpha oscillations with age (***Figure 6 – figure supplement 1***).

Slow, delta and alpha oscillations were the dominant frequency components which evolved in the level of frontal coherence observed across age. Peak frontal coherence was evaluated for slow-delta and alpha frequencies across age (***Figure 7***). Frontal slow-delta oscillations were highly coherent from birth and steadily decreased with age until around 8 months old (***Figure 7A & B***). Frontal alpha oscillations were incoherent from birth, until 10 months of age (after appearing in the spectrogram at 3-4 months) and gradually increased coherence with increasing age (***Figure 7C***).

**Figure 7.**
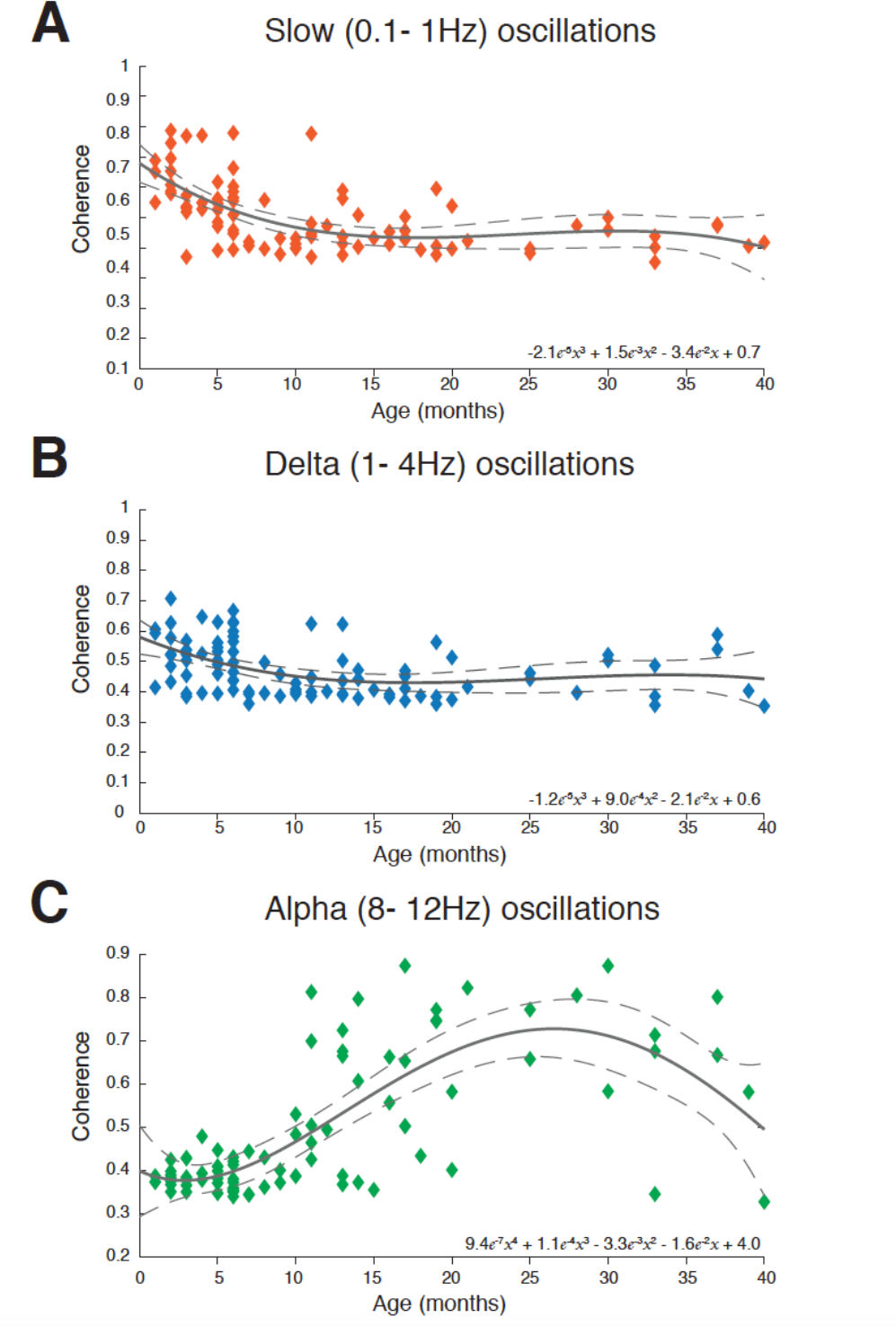
Trends in frontal slow, delta and alpha coherence with age from 0 to 3 yrs. Frontal coherence in the (A) slow (0.1 to 1Hz), (B) delta (1-4Hz), and (C) alpha (8-12Hz) frequency bands for each subject during sevoflurane-maintained surgical anesthesia, plotted as a function of age. Slow (0.1-1Hz), and delta (1-4Hz) coherence are present in subjects from birth until around 8 months old. Alpha coherence appears to emerge around 10 months of age and gradually increases in 20 months where it persists to 3 years of age. Solid line represents a third or fourth-degree polynomial regression model describing the relationship between age and EEG power; dashed lines represent the 95^th^ percentile confidence boundaries of this regression model. F7-Fp1 vs. F8-Fp2 used with nearest neighbor Laplacian referencing. Epoch size of 5 minutes is used.

#### Global coherence is weak across all frequencies in the first 9 months of age; frontal alpha coherence emerges at around 10 months of age

Global coherence analysis characterizes coordinated activity across multiple channels of EEG as a function of frequency. Global coherence is the fraction of variance at a given frequency across all EEG channels that is explained by the first eigenvector of the cross-spectral matrix. Topographic maps of coherence were computed for slow-delta, theta, alpha, and beta frequency bands in each age-group (***Figure 8***). Despite frontally coherent slow-delta oscillations in infants from 0 to ~8 months of age, global coherence was weak. In contrast, frontally coherent alpha oscillations were present in infants from 10 months of age, with a global coherence projection weight of around 0.3 (***Figure 8***). The global coherence in the slow and delta frequency ranges is consistent with the lack of slow oscillation coherence observed in adults (Cimenser et al., 2011; Lewis et al., 2012; Purdon et al., 2013). Furthermore, coherence in the alpha-frequency range is similar to the highly coordinated, coherent frontal alpha activity observed in adults during *propofol* general anesthesia (Cimenser et al., 2011; Purdon et al., 2013).

**Figure 8.**
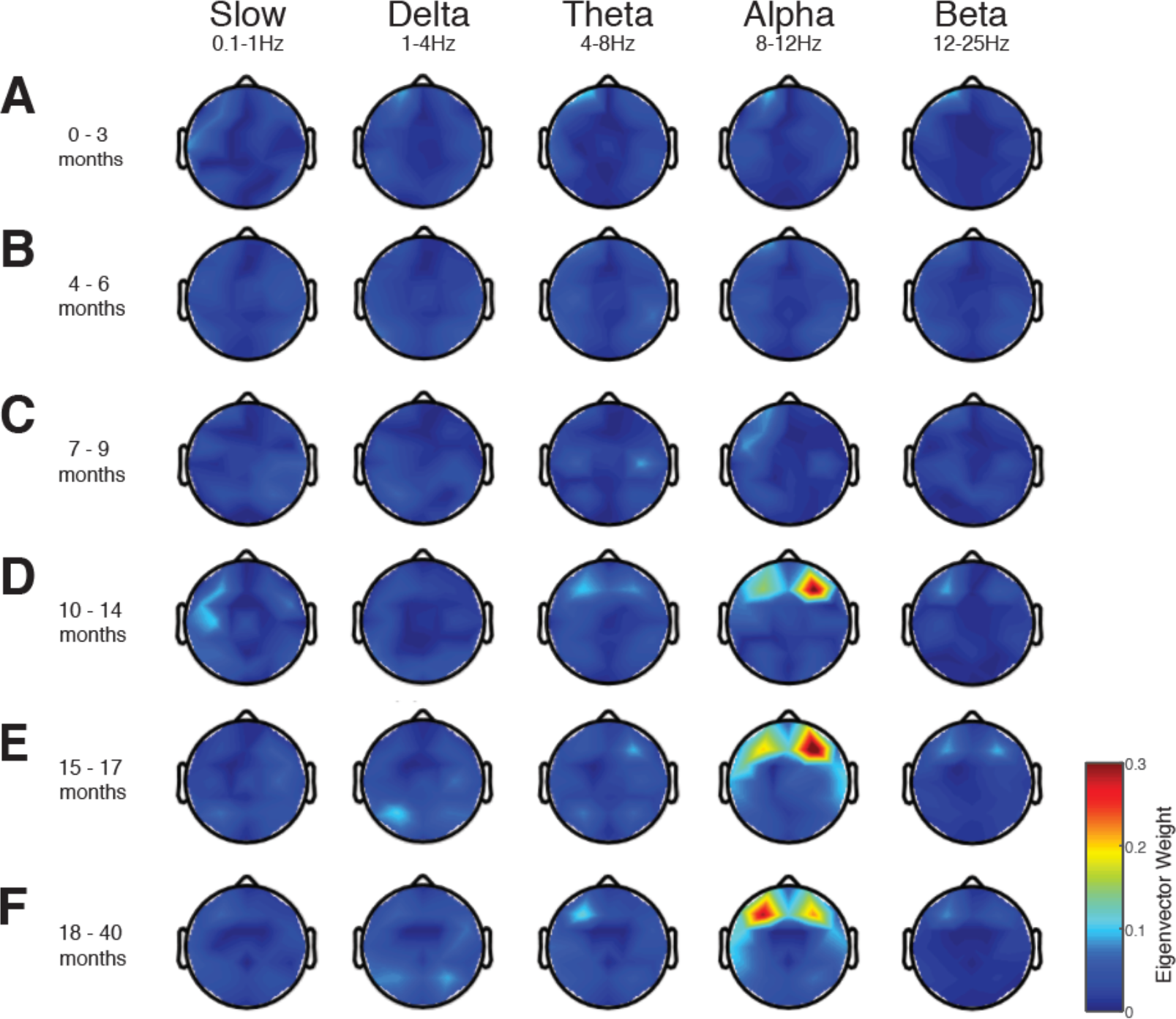
Global coherence during sevoflurane-maintained surgical-anesthesia in children from 0 to 3 yrs. Topographic EEG maps detailing group-averaged coherence for each EEG frequency band in children aged (A) 0-3 months (n=17), (B) 4-6 months (n=23), (C) 7-9 months (n=7), (D) 10-14 months (n=18), (E) 15-17 months (n=7), and (F), and 18-40 months (n=19). Spatially coherent alpha oscillations that are concentrated in the frontal channels emerge at 10-14 months of age. Global coherence is low across all remaining frequencies in all ages during surgical anesthesia. This analysis suggests that there is an age-varying change in spatially coherent alpha oscillations during late infancy phase of brain development.

## Discussion

### Summary of findings

In this study, we found age-dependent changes in the EEG during exposure to sevoflurane general anesthesia. Specifically, we found that: (1) slow-delta oscillations are present in all ages (***Figure 2***); (2) theta and alpha oscillations emerge around 4 months (***Figure 2***); (3) alpha oscillations increase in power from 4 to 10 months, where they peak and remain at a sustained power level at older ages (***Figure 3***); (4) frontal alpha oscillation predominance emerges around 7 months, and continue to exhibit a dominant anterior spatial distribution at older ages (***Figures 4 & 5***); (5) frontal slow oscillations are coherent from birth until around 6 months when they start to become incoherent (***Figures 6 & 7***); and (6) frontal alpha oscillations become coherent around 10 months and persist in older ages (***Figures 6-8)***. These age-related changes in the EEG likely relate to changes in neurodevelopmental processes that occur in early childhood, including synaptogenesis, axonal growth, neural pruning, myelination, and the maturation of thalamocortical and cortico-cortical functional connectivity (Andersen, 2003; Tau & Peterson, 2010; Marín, 2016; Verriotis et al., 2016; Cao et al., 2017). These unique data provide putative markers of the postnatal maturation GABAergic-driven circuits in the human infant brain (***Figure 9***).

#### Studies of EEG dynamics in adults under general anesthesia

Monitoring brain activity in response to general anesthesia has been extensively characterized in adults. Specifically, the brain states induced by general anesthetic drugs have been well-studied through detailed characterizations of behavior and consciousness at varying drug levels in adult human volunteers or in surgical patients (Gugino et al., 2001; Mhuircheartaigh et al., 2010; Ku et al., 2011; Purdon et al., 2013; Akeju et al., 2014; Mhuircheartaigh et al., 2013; Blain-Moraes et al., 2015; Barttfeld et al., 2015; Akeju et al., 2016; Hight et al., 2017; Pavone et al., 2017).

A dose-dependent effect on frontal predominance of alpha oscillations under deep sedation with sevoflurane occurs in healthy adult volunteers (Gugino et al., 2001). More recently, high-density EEG demonstrated that unconsciousness induced by sevoflurane or propofol is associated with a functional disconnection in the alpha oscillations between the anterior and posterior structures (Blain-Moraes et al., 2015; Cimenser et al., 2011; Pavone et al., 2017; Purdon et al., 2013). Frontal and thalamocortical information processing are thought to play a major role in the generation of consciousness, and impairment of this processing might be an important neurophysiologic mechanism of the action of general anesthetics.

#### Studies of EEG dynamics of the infant brain under general anesthesia

We observed that EEG functional networks increase in density over development, and are established in an “adult-like” manner for sevoflurane-associated anesthesia by late-infancy. Age-related differences in EEG signatures have been shown in previous work in infants and toddlers. Davidson *et al* monitored 8-leads of EEG in 17 infants (0-6 months), 19 toddlers (6 months to 2 years) and 21 children (2 to 10 years) during emergence from anesthesia. They found that power –limited to 2 to 20 Hz-was lower during anesthesia in infants compared to older children, and that power was similar during anesthesia in toddlers and children (Davidson et al., 2008). Hayashi *et al* conducted a retrospective study on EEG data collected using a 2-lead EEG from 62 children ranging from 1 day to 2 years of age who received sevoflurane anesthesia. They found that the spectral edge frequency was low even under light anesthesia and changes with deeper anesthesia were small in infants younger than 3 months of age (Hayashi et al., 2012). Sury *et al* studied 20 infants at 1 week to 10 months of age during sevoflurane anesthesia using a 4-lead EEG and analyzed spectral power between 1 and 28Hz. They found that infants 3 months and older had greater power in the 2-20Hz range during maintenance compared to younger infants (Sury et al., 2014).

Akeju *et al* retrospectively characterized frontal EEG data using a 4-lead Sedline brain function monitor (electrode positions: FP1, FP2, F7 and F8) in 54 subjects ranging in age from 0.4 to 28 years (of these, 4 subjects were <1 year of age). They analyzed frontal power spectra and coherence changes with age. They observed that frontal slow oscillations were present from birth to 28 years, and frontal alpha oscillations emerged around 1 year of age. They also found age-dependent changes in coherence where slow frequencies were transiently coherent – from birth to 1 year of age – whereas alpha oscillations were coherent around 1 year of age and older, reflecting a pattern of power and coherence similar to adults (Akeju et al., 2015).

We recently characterized temporal changes in the EEG using multi-electrode recordings in 36 infants 0 to 6 months old when awake, and during maintenance of and emergence from sevoflurane (Cornelissen et al., 2015). In agreement with the current data, we found that slow oscillations were present and coherent in all infants, and alpha oscillations emerged around 3 to 4 months of age and remained incoherent. The coherent frontal alpha oscillations seen in adults are thought to be a GABA-associated thalamocortical oscillation (Ching et al., 2010). Age-dependent changes in coherence might reflect functional changes in GABA-associated frontal thalamocortical circuits (***Figure 9***).

**Figure 9.**
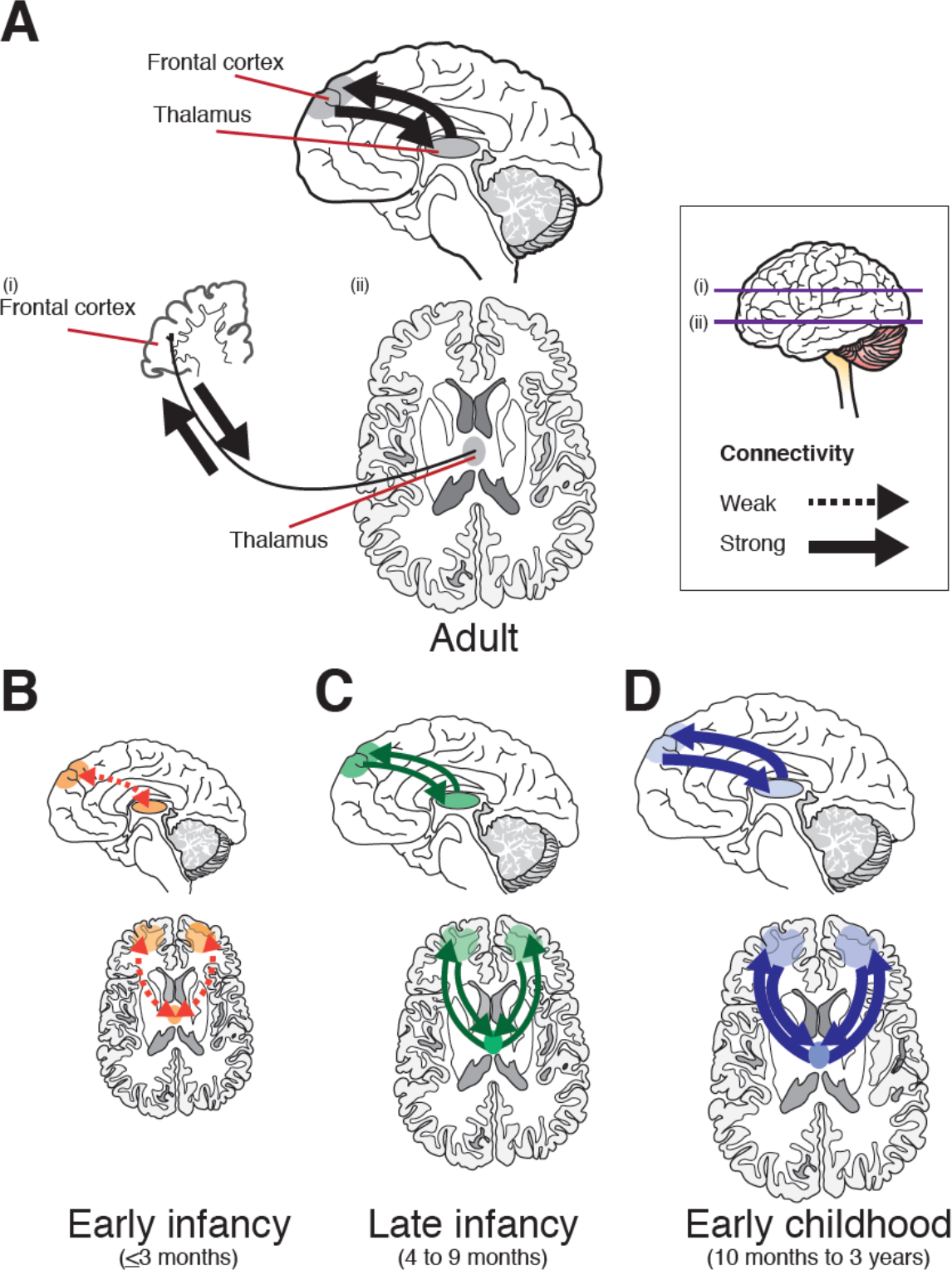
Hypothetical model of age-varying changes in thalamocortical and cortico-thalamic connectivity during sevoflurane general anesthesia. (A) In adults, sevoflurane and propofol general anesthesia are characterized by anteriorization of alpha oscillations which are driven by frontal thalamocortical activity. (B) In infants 0 to 3 months of age, frontal alpha oscillations are absent and this is likely due to weak connectivity of projections from the thalamus to the frontal cortices and from the cortices to the thalamus. (C) In infants, from 4 to 9 months of age, alpha oscillations are observed and located over the frontal cortex. The appearance of frontal alpha oscillations is thought to be driven by maturation of thalamocortical and cortico-thalamic connections. Alpha oscillations remain incoherent across the left and right frontal hemispheres at these ages and this may be due to immaturity of the corpus callosum, which serves to join the two hemispheres of the brain, and while in place from birth undergoes rapid maturation over the first 9 months of life. (D) Towards the end of infancy and into early childhood, children aged between 10 months and 3 years show highly synchronous and robust frontal alpha oscillations that mimic patterns of neural activity that are similar to that of the adult under sevoflurane or propofol general anesthesia and likely reflect highly function thalamocortical and cortico-thalamic connectivity. Regions of interest shown in gray (Adult), orange (0 to 3months of age), green (4 to 9 months of age), and blue (10 months to 3 years of age).

Few studies have evaluated spatial changes in the EEG during anesthesia in young chidren. Lo *et al* characterized spatial EEG patterns in 37 children 22 days to 3.6 years who received sevoflurane or isoflurane anesthesia using 128 leads. They found spatial differences in the power of EEG frequency bands –ranging from 1 to 50Hz-over the frontal cortex compared to the occipital cortex at high concentrations of the two agents. However, none of the analyses were stratified by age (Lo et al., 2009).

#### Postnatal development of brain structure and function

Age-related changes we see in the EEG are consistent with the neurobiology and neurophysiology of early development. The dramatic maturation of different cortical rhythms during the first year of life is likely due to synaptogenesis, dendrite elaboration, myelination, white matter tract development and continued sub-plate growth. In this study, we observe that at younger ages, <3 months of age, before thalamocortical and cortico-thalamic connections are established, slow oscillations are dominant due to cortical hyperpolarization. At 3 to 9 months, thalamocortical and cortico-thalamic connections are weakly established with minimal to no-coherence in the alpha range observed. At older ages, ≥10 months of age, robust thalamocortical and cortico-thalamocortical connections are present and consequently highly coherent alpha oscillations are observed (Figure 9).

Both cortico-cortical and thalamocortical connections are formed during the second half of gestation, and these connections play a role in synchrony and inter hemispheric connectivity (Fransson et al., 2007; Fransson et al., 2011; Kostović & Jovanov-Milosević, 2006; Kostović & Judas, 2010). At birth, a ‘*rich-club’* organization with specific cortical *hubs* connected to each other is already in place, and is thought to provide a foundation for postnatal development of functional connectivity networks (Ball et al., 2014).

The brain grows considerably in size during the first 3 years. While the left and right hemispheres develop at different rates, with different postnatal onset times, metabolic changes associated with rapid synaptogenesis and axonal growth are observed during the first three to six months of life (Blüml et al., 2013; Kehrer & Schöning, 2009). Structural MRI studies indicate the corpus callosum, which serves to join the left and right hemispheres of the brain, is uniformly thin during the neonatal period, and undergoes rapid growth spurts from 3 months onwards until it begins to have an adult-like appearance by about 9 months of age (Barkovich & Kjos, 1988; Provenzale et al., 2012). Myelin is already present during the first year in all associated regions but maturation is ongoing -with the frontal cortex being one of the last to fully myelinate (Brody et al., 1987; Kinney et al., 1988). By 2 years of age, cortical thickness reaches approximately 97% of adult values, whereas cortical surface area only reaches 69% of adult values (Li, et al., 2015). By 3 years of age, neuron and glial counts remain stationary in number, and myelination (with the exception of the reticular formation, for example) is approximately 90% of adult values. Frontal areas and cortico-cortical connections continue to mature until puberty (Emerson et al., 2016; Marín, 2016; Thatcher, 1992; Thatcher et al., 1987; Thatcher et al., 2008). Significant advances in motor, cognitive, and linguistic network development occur over this time, as well as a number of higher order cognitive functions such as face recognition (Pascalis et al., 2002), and self-awareness (Kristen-Antonow et al., 2015).

#### Clinical implications for brain monitoring under anesthesia

Critical periods of brain development are strongly influenced by the maturation of GABAergic interneurons, and represent a major milestone in the development of neural circuits. Critical period plasticity is diverse in onset and duration, and occur at different times depending on the function. Animal studies show these critical periods can be prematurely opened, kept open or prematurely closed by exposure to GABA-acting compounds such as benzodiazepines (Hensch, 2005).

Preterm birth, and neurodevelopmental disorders such as autism spectrum disorder, are associated with lower alpha oscillatory power during general anesthesia compared to controls (Poorun et al., 2015; Walsh et al., 2016). Our results encourage further examinations of large-scale functional brain network in “typically developing infants”, as well as in infant exposed to repeated anesthesia or prolonged sedation with GABAergic agents, and in neurodevelopmental pathologies involving altered excitatory-inhibitory balance such as Rett syndrome or Fragile X syndrome.

Concerning results from animal studies suggest that sustained exposure to GABA_A_ receptor agonists and NMDA antagonists in the neonatal and infant period accelerates neurodegenerative mechanisms resulting in impaired learning and memory skills in later life (Vutskits & Xie, 2016). In human studies, the evidence is less clear and remains hotly debated as accurate translation of animal behavior to human outcomes is challenging. Some retrospective studies suggest repeated exposure to general anesthesia in early life (<2 years) may increase the risk for adverse long-term neurodevelopmental, while more recent prospective studies suggest otherwise (Vutskits & Xie, 2016). Coordinating timing and duration of exposure to general anesthetics to avoid at critical periods responsible for learning and memory acquisition may significantly reduce vulnerability to negative outcomes (Andropoulos & Greene, 2017). ln understanding the EEG correlates of sevoflurane-induced unconsciousness, we may be able to optimize clinical monitoring approaches and eliminate risk of anesthetic-induced neurotoxicity in the developing brain.

#### Study limitations and constraints

Anesthetic management was administered according to the anesthesiologist’s discretion, rather than in a controlled, titrated fashion. It is possible that the observed EEG features could have been confounded by systematic age-related differences in drug administration. To prevent this, analysis epochs were identified at a period of surgical anesthesia and reflect a brain state consistent with anesthesia-induced unconsciousness across the sample population. The spectral and coherence effects are unlikely to be the result of over anesthetizing younger subjects, and are more likely age-related differences in underlying neurophysiology. Although challenging to pursue, there is a need for more detailed EEG studies featuring controlled drug administration and pharmacokinetic measurements in children.

## Conclusions

Our findings are consistent with recent work over smaller and more sparsely sampled age ranges. We show that general anesthetics that primarily act on the GABAergic system can provide windows of cortical maturation and integration through robust neurophysiological signatures throughout development.

## Methods and Materials

### Study Design

The objective of this study was to evaluate the effect of postnatal age on electroencephalographic (EEG) activity during sevoflurane general anesthesia in children 0-3 years old. We recorded multichannel EEGs during administration of sevoflurane general anesthesia for elective surgery (per clinical protocol). End-tidal anesthetic gas volume and video recordings of behavioral activity were time-locked to the EEG recording. The spatial and temporal properties of the pediatric EEG were evaluated during a sevoflurane-maintained surgical state of anesthesia.

#### Participants

Children who were scheduled for an elective surgical procedure were recruited from the pre-operative clinic at Boston Children’s Hospital from December 2011 to August 2016. Eligibility criteria consisted of children aged 0 to 3 years, who required surgery below the neck. All children were clinically stable on the day of study and American Society of Anesthesiologists’ physical status I or II. Exclusion criteria were (1) born with congenital malformations or other genetic conditions thought to influence brain development, (2) diagnosed with a neurological or cardiovascular disorder, or (3) born at <32 weeks post-menstrual age.

This study was approved by the Boston Children’s Hospital Institutional Review Board (Protocol Number P000003544) and classified as a ‘*no more than Minimal Risk’* study. Informed written consent was obtained from the parents or legal guardians before each study. The study conformed to the Declaration of Helsinki and Good Clinical Practice guidelines.

A total of 106 EEG recordings were performed. Data are presented from 91 subjects (0-3 months, n=17; 4-6 months, n=23; 7-9 months, n=7; 10-14 months, n=18; 15-17 months, n=7; and 18-40 months, n=19) in the analysis (***Figure 1-figure supplement 1***). Summary demographics of subjects included in the analysis are given in ***Table 1***.

#### Anesthetic Management

Each patient received anesthesia induced with sevoflurane alone, or a combination of sevoflurane and nitrous oxide. Nitrous oxide was added at the discretion of the anesthesiologist (67 subjects). Nitrous oxide was discontinued after placement of an endotracheal tube or laryngeal mask. Endotracheal tube airway was used in 62 subjects, laryngeal mask airway was used in 24 subjects, and face mask was used in 5 subjects. Propofol bolus was used to facilitate tracheal intubation or suppress motor reflexes in 44 subjects [median dose: 2.1mg/kg (95% CI: 1.7-2.4 mg/kg)]. Seven subjects were prescribed midazolam premedication on the day of surgery.

Epochs used for analysis were comprised of sevoflurane administration with air and oxygen, titrated to clinical signs; end-tidal sevoflurane concentration was adjusted per the anesthesiologist’s impression of clinical need, not a pre-set end-tidal sevoflurane concentration (***Table 1***).

## Data Acquisition

All subjects were in the supine position throughout the study. Each subject was studied once.

### EEG Recording

An EEG cap was used to record EEG activity (WaveGuard EEG cap, Advanced NeuroTechnology, Enschede, Netherlands). 33- or 41-recording electrodes were positioned per the modified international 10/20 electrode placement system. For 33-channel recording, electrodes were positioned at FPz, FP1, FP2, F3, F4, F7, F8, FC1, FC2, FC5, FC6, Cz, CPz, C3, C4, CP1, CP2, CP5, CP6, Pz, P3, P4, P7, P8, T7, T8, M1, M2, POz, Oz, O1 and O2 (***Figure 1***). For 41-channel recording, additional electrodes were positioned at AF7, AF8, PO7, PO8, FT7, FT8, TP7 and TP8. Reference and ground electrodes were located at Fz and AFz respectively. The impedance of the electrode-skin interface was kept to a minimum by massaging the skin with an EEG prepping gel (Nu-Prep gel, DO Weaver & Co., CO, USA), and conductive EEG gel was used to optimize contact with the electrodes (Onestep-Clear gel, H^+^H Medical Devices, Dulmen, Germany).

EEG activity from 0-500 Hz was recorded with an Xltek EEG recording system (EMU40EX, Natus Medical Inc., Ontario, Canada). Signals were digitized at a sampling rate of 1024Hz (or 256Hz in 5 cases), and a resolution of 16-bit.

#### Clinical Data Collection

Demographics and clinical information, including age, gender, surgical procedure, anesthetic management, were collected from the electronic medical records and from the in-house Anesthesia Information Management System (AIMS). End-tidal sevoflurane, oxygen, and nitrous oxide concentrations were downloaded from the anesthetic monitoring device (Dräger Apollo, Dräger Medical Inc., Telford, PA) to a recording computer in real-time using ixTrend software (ixcellence, Wildau, Germany). Signals were recorded at a 1 Hz sampling rate.

#### EEG Analysis

##### Data Preprocessing

Preprocessing was carried out with Natus Neuroworks (Natus Medical Inc., Ontario, Canada) and MATLAB (MathWorks, Natick, MA). Ear electrodes (M1 and M2) were excluded from the final analysis due to poor surface-to-skin contact for most subjects.

EEG signals were re-montaged to a nearest neighbor Laplacian reference using distances along the scalp surface to weight neighboring electrode contributions. We applied an anti-aliasing filter of 80 Hz and down-sampled the EEG data to 256 Hz.

For each subject, a period of maintenance of general anesthesia adequate for surgery and where end-tidal sevoflurane concentration was maintained at a steady concentration (+/-0.1 %) was identified in the EEG. Within this segment, a 5-minute epoch was selected from ‘artefact-free’ EEG, where motion or electrocautery artefacts were not present in the EEG. Channels with noise or artefacts were excluded from the analysis by visual inspection. EEG data epochs were analyzed with a median time after the start of the surgical procedure of 15 min (95% CI: 10 – 23 min). Two authors (L.C., J.M.L) visually inspected all EEG data for each subject and manually selected data ‘artefact-free’ EEG segments for analysis.

##### Time-frequency Analysis

Spectral analysis of activity was performed with multitaper methods using the Chronux toolbox (http://chronux.org) (Bokil et al., 2010). Multitaper parameters were set using window lengths of T=2 seconds with a 1.9 second overlap; time-bandwidth product, TW = 2; and number of tapers, K=3. The spectrum of frequencies over time within the 0-40Hz bandwidth were plotted for individual electrodes in each subject.

Subjects were divided into groups per postnatal age: (i) 0-3 months, (ii) 4-6 months, (iii) 7-9 months, (iv) 10-14 months, (v) 15-17 months, and (vi) 18-40 months of age. First, group-averaged spectrograms were computed by taking the median power across subjects at each time and frequency at the electrode of interest for each postnatal age cohort (i.e. ***Figure 2 – figure supplement 1A***). Second, group-averaged spectra were computed by taking the median power across subjects at each frequency across the entire 5-minute epoch, and then the median (95% CI) power at each frequency was calculated for each postnatal age group (i.e. ***Figure 2 – figure supplement 1B***). To identify age-varying changes in the frontal power spectra, we computed an age-varying spectrogram using an overlapping (1-month) moving window spanning a 3-month range. To obtain smoothed spectrograms, we applied a Kalman filter algorithm and a fixed interval smoothing algorithm to the age-varying spectrogram (i.e. ***Figure 2***) (Mendel, 1995).

To identify the topographic distribution for specific frequency bands, we first took the mean power spectra of group-averaged spectrograms across the entire 5-minute epoch for each electrode. Then, we averaged the mean power spectra over each EEG frequency band of interest for each postnatal age group. Scalp EEG frequency band power plots were performed using 3D interpolation of the electrode montage with the topoplot function in EEGLab (Delorme & Makeig, 2004).

#### Coherence analysis

Coherence analysis was performed using custom-written MATLAB code (MathWorks Inc., Natick, MA) (Cornelissen et al., 2015). Coherence quantifies the degree of correlation between two signals at a given frequency. It is equivalent to a correlation coefficient indexed by frequency: a coherence of 1 indicates that two signals are perfectly correlated at that frequency, while a coherence of 0 indicates that the two signals are uncorrelated at that frequency. The coherence between signals *i* and *j* is given by:

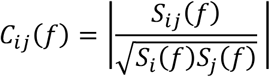

where *S_ij_* is the cross-spectrum between signals *x_i_*(t) and *x_j_*(t), *S_i_* is the power spectrum of *x_i_*(t), and *S_j_* is the power spectrum of *x_j_*(t).

*For frontal coherence analysis*, we used raw EEG data recorded using a common reference (Fz) to avoid the distortion by referencing. Using the raw EEG data, we estimated coherence between bipolar left frontal signal [F7-Fp1] and bipolar right frontal signal [F8-Fp2], using multitaper methods with TW = 4, K = 7, and t = 4s with a 3 second overlap based on analysis of adult EEG (Akeju et al., 2014). A coherogram graphically illustrates coherence for a range of frequencies plotted across time. To compute the group-averaged coherogram, we took the median coherence across subjects at each time point and frequency. To identify age-varying changes in frontal coherence, we computed an age-varying coherogram using an overlapping (1-month) moving window spanning a 3-month range and applied the Kalman and smoothing filter used in the spectral analysis (i.e. ***Figure 6***).

*For global coherence analysis*, we divided each segment into non-overlapping 2 second windows and computed the cross-spectral matrix. To remove noise artefact. The median over 10 windows of the real and imaginary parts of each entry in the cross-spectral matrix was taken (Wong et al., 2011). Then, we performed an eigenvalue decomposition analysis of the cross-spectral matrix at each frequency. The cross-spectral matrix at each frequency can be factorized as:

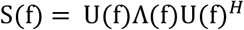

where U^H^ is the complex conjugate transpose of U and a unitary matrix whose ith column is the eigenvector u_i_ of S, and Ʌ is the diagonal matrix whose diagonal elements are the corresponding eigenvalues,Ʌ_ii_ =λ_i_. The global coherence is the ratio of the largest eigenvalue to the sum of eigenvalues (Cimenser et al., 2011):

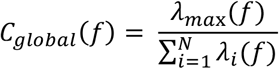

When the largest eigenvalue is large compared with the remaining ones, the global coherence is close to 1. We computed the global coherence at each frequency using 5-minute epochs for MOSSA. To determine group-averaged global coherence, we took a median across subjects in each postnatal age group. We refer to the eigenvector u_max_ corresponding to the largest eigenvalueλ_max_ at a given frequency as the principal mode of oscillation for that frequency and the coherence of electrode sites was obtained by the absolute square of the eigenvector as previously applied in adult EEG studies (Purdon et al., 2013). This means that the eigenvector described a coherent spatial distribution. Spatial coherence at each frequency band was computed by taking an average across the frequency range at each electrode. Group-averaged scalp coherence distribution was computed by taking the median across subjects and plotting with the topoplot function in EEGlab.

### Statistical Analysis

Data were tested for normality using a D-Agostino normality test. Data are shown as median (95% CI of the population median) unless otherwise stated. Analyses were performed using R v3.4.3 (“*You Stupid Darkness*”) with *MedOr* (for confidence interval of population medians analysis), *moments* (for normality testing), and *tidyverse* packages (http://www.R-project.org) (R Core Team, 2013), and custom-written MATLAB code [MathWorks Inc., Natick, MA; (Cornelissen et al., 2015)].

We used the *polyfit* function in MATLAB (MathWorks, Natick, MA) to obtain the best-fit regression to describe the relationship between age and power, or age and coherence and the *polyconf* function in MATLAB (MathWorks, Natick, MA) to obtain the confidence intervals. We used a paired-subject bootstrapping algorithm to compare spectral estimates between electrode locations (frontal vs occipital), as implemented in the Chronux toolbox (Kirch & Politis, 2011). We drew bootstrap samples from the data set with replacement and estimated the best-fit regression for the bootstrap samples using the *polyfit* function for each electrode location. Then we took the difference between regression functions at specific electrode locations (using paired comparisons). We repeated this process 10,000 times and calculated the 95% CI for the difference between two regression functions to test for significant differences.

### Use of Previously Published Datasets

Previous papers describing EEG characteristics during general anaesthesia on selected subjects are reported elsewhere: specifically, on anesthesia-associated EEG spectral and coherence patterns in infants from 0-6 months (n=36) (Cornelissen et al., 2015), and on EEG discontinuity (profoundly suppressed EEG activity) during deep levels of anaesthesia in children from 0 to 3 yrs (n=68) (Cornelissen et al., 2017).

### Code

Custom-written MATLAB code (with simulated data) for computing multitaper spectra, global coherence and bootstrap CIs is available through our previous article on infants in *eLife* (Cornelissen et al., 2015).

## Acknowledgments

We thank the Pre-operative and Operating Room staff at Boston Children’s Hospital for their assistance during these studies, as well as the families who took part in the study. We also thank Dr. Ann-Marie Bergin in the Department of Neurology for reviewing all EEG recordings for potential incidental findings.

## Additional information

### Funding

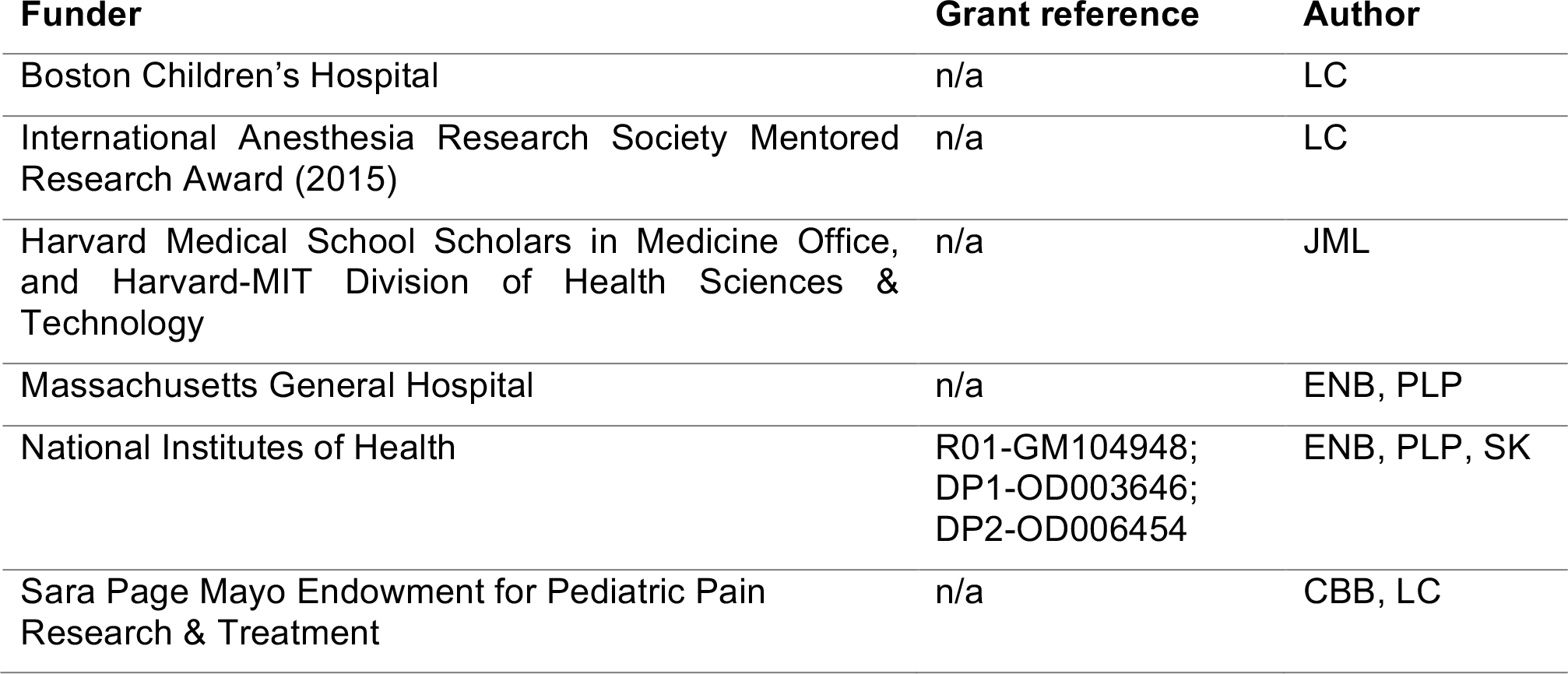

### Author contributions

The work presented here was undertaken in collaboration between all authors. LC and CBB designed the study. LC and JL acquired the data. SK, PLP and ENB designed the computational analysis. SK, LC and JL carried out the data analysis. All authors contributed to data analysis and data interpretation. LC, PLP, CBB and ENB wrote the initial manuscript; all authors edited and revised the manuscript. All authors approved the final version of this manuscript.

### Author ORCIDs (if available)

Laura Cornelissen, http://orcid.org/0000-0001-85790870

Seong-Eun Kim: http://orcid.org/0000-0002-4518-4208

Patrick Purdon, http://orcid.org/0000-0001-5651-5060

Emery Brown, http://orcid.org/0000-0003-2668-7819

### Ethics

Human subjects: This study was approved by the Boston Children’s Hospital Institutional Review Board (Protocol Number P000003544) and classified as a ‘*no more than Minimal Risk’* study. Informed written consent was obtained from the parents or legal guardians before each study. The study conformed to the Declaration of Helsinki and Good Clinical Practice guidelines.

## Additional files

### Supplementary files

#### Supplementary File 1

Figure 1 – supplement 1.

Study profile

#### Supplementary File 2

Figure 2 – figure supplement 1.

Frontal EEG spectral properties across age-groups

#### Supplementary File 3

Figure 4 – figure supplement 1.

Frontal coherence properties during sevoflurane-maintained surgical anesthesia across age-groups

#### Supplementary File 4

Figure 6 – figure supplement 1.

Frontal coherence properties during sevoflurane-maintained surgical anesthesia across age-groups

